# Riboglow: a multicolor riboswitch-based platform for live cell imaging of mRNA and small non-coding RNA in mammalian cells

**DOI:** 10.1101/199240

**Authors:** Esther Braselmann, Aleksandra J. Wierzba, Jacob T. Polaski, Mikołaj Chromiński, Zachariah E. Holmes, Sheng-Ting Hung, Dilara Batan, Joshua R. Wheeler, Roy Parker, Ralph Jimenez, Dorota Gryko, Robert T. Batey, Amy E. Palmer

## Abstract

RNAs directly regulate a vast array of critical cellular processes, emphasizing the need for robust approaches to fluorescently tag and track RNAs in living cells. Here, we develop an RNA imaging platform using the cobalamin riboswitch as an RNA tag and a series of probes containing cobalamin as a fluorescence quencher. This highly modular ‘Riboglow’ platform leverages different color fluorescent dyes, linkers and riboswitch RNA tags to elicit fluorescent turn-on upon binding RNA. We demonstrate the ability of two different Riboglow probes to track mRNA and small non-coding U RNA in live mammalian cells. A direct side-by-side comparison revealed that Riboglow outperformed the dye binding aptamer Broccoli and performed on par with the current gold standard RNA imaging system, the MS2-fluorescent protein system, while featuring a much smaller RNA tag. Together, the versatility of the Riboglow platform and ability to track diverse RNAs suggest broad applicability for a variety of imaging approaches.

**Graphical abstract:** 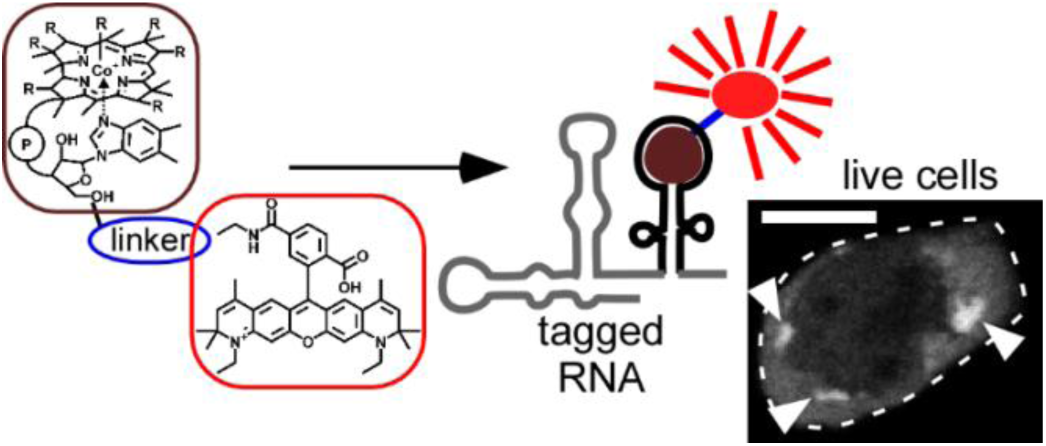

## Introduction

The complex spatiotemporal dynamics of messenger RNAs (mRNAs) and non-coding RNAs (ncRNAs) affect virtually all aspects of cellular function. These RNAs associate with a large group of RNA binding proteins that dynamically modulate RNA localization and function^1,2^ and such interactions govern mRNA processing, export from the nucleus, and assembly into translationally competent messages, as well as association into large macromolecular granules that are not translationally active, including processing bodies (P-bodies) and stress granules (SGs)^3–6^. Similarly, uridine-rich small nuclear RNAs (U snRNAs, the RNA components of the spliceosome)^7^ dynamically associate with protein components to comprise the functional spliceosomal complex in the nucleus^8^. During stress, such as nutrient deprivation or bacterial infection, U snRNAs along with the splicing machinery can be transiently sequestered in cytosolic foci called U-bodies^7^. Given the intricate connection between RNA localization, dynamics and function, there has been a strong push to develop tools for visualization of RNA in live cells to elucidate mechanisms underlying dynamics of the mRNA and ncRNA life-cycle.

While there is a broad spectrum of tools to fluorescently tag proteins in live cells, fewer approaches for live cell imaging of RNA exist. The most common system employs multimer RNA tags that bind an RNA-binding protein (MS2 or PP7 coat protein) fused to a fluorescent protein (FP)^9–11^. The tag is genetically fused to an RNA of interest and binding of MS2-FP concentrates the fluorescence signal on the RNA. However, one limitation of this approach is that many copies of the hairpin RNA tag are required to enhance fluorescence contrast, and the large size of the RNA tag (Table 1) bound to FP complexes can perturb localization, dynamics and processing^12^ of the RNA. Still, this system is the gold standard in live cell RNA imaging as it has been used successfully to interrogate mRNA dynamics over time at the single molecule level^9,10,13–15^. An alternative approach involves fluorogenic dye-binding aptamers that give rise to a turn-on fluorescence signal when the dye binds the aptamer^16-20^. While numerous proof-of-principle aptamers have been developed^21^, only the Spinach^22^, Broccoli^23^ and Mango^24,25^ aptamers have been used in live mammalian cells. These dye-binding aptamers have been used to visualize highly expressed RNA polymerase III-dependent transcripts such as 5*S* and U6 RNA^25–27^. However, there are no reports of dye-binding aptamers being used to detect RNA polymerase-II dependent transcripts such as mRNAs, snRNAs, or microRNAs. These tools demonstrate the power and potential of RNA imaging to reveal spatiotemporal dynamics of RNA function. But, they also highlight the need for robust and complementary approaches for fluorescently tagging and visualizing diverse types of RNA in live mammalian cells with small molecular probes.

**Table 1.** Comparison of length and fluorophore properties for RNA tags used for live cell RNA fluorescence microscopy in this study.

| Name of tag | Length of RNA tag<br>(nucleotides) | number of dye / FP<br>molecules per tag | Molecular weight (kDa) of<br>fluorophore |
| --- | --- | --- | --- |
| (A<sub>T</sub>)1x | 81 | 1 | 2.3 (Cbl-5xPEG-ATTO 590) |
| (A)1x | 103 | 1 | 2.3 (Cbl-5xPEG-ATTO 590) |
| (A)2x | 212 | 2 | 2.3 (Cbl-5xPEG-ATTO 590) |
| (A)3x | 321 | 3 | 2.3 (Cbl-5xPEG-ATTO 590) |
| (A)4x | 430 | 4 | 2.3 (Cbl-5xPEG-ATTO 590) |
| 1x MS2 SL | 61 | 2 | ~88 (MS2-GFP homo dimer) |
| 2x MS2 SL | 102 | 4 | ~176 |
| 4x MS2 SL | 210 | 8 | ~352 |
| 24x MS2 SL | 1354 | 48 | ~2,112 |
| 2xdBroccoli | 234 | 4 | 0.3 (DFHBI-1T) |

## Results

### Design of RNA-binding probes with fluorescence quenching properties

Here, we introduce an orthogonal approach for fluorescent tagging of RNA in live cells using a bacterial riboswitch as the RNA tag and a series of small molecular probes that increase in fluorescence upon binding the RNA. We took advantage of the robust folding of bacterial riboswitches in different genetic contexts in cells^28,29^, while exploiting specific binding of the riboswitch RNA to its natural ligand, cobalamin (Cbl)^30,31^. Cbl functions as an efficient fluorescence quencher when covalently coupled to a synthetic fluorophore^32–34^. We postulated that binding of Cbl to an RNA riboswitch could sterically separate the Cbl quencher and covalently coupled fluorophore resulting in fluorescence turn-on of a Cbl-fluorophore probe (Fig. 1a), similar to fluorophore-quencher systems used for RNA tagging *in vitro* and in bacteria^18,19^. The Cbl riboswitch consists of the aptamer (the Cbl-binding domain) and the expression platform. Variants of both the full-length riboswitch sequence and the shorter aptamer domain were used (Supplementary Fig. 1), collectively referred to as the ‘riboswitch RNA tag’ in this study. The crystal structure of Cbl bound to the aptamer revealed that the Cbl 5’-hydroxyl group is accessible even in the RNA-bound state^30^ (Fig. 1b), leading us to choose this position for conjugation with fluorophores via the copper catalyzed alkyne-azide cycloaddition reaction^35,36^ to explore different conformational and photochemical properties of the probe (Fig. 1c).

**Figure 1:**
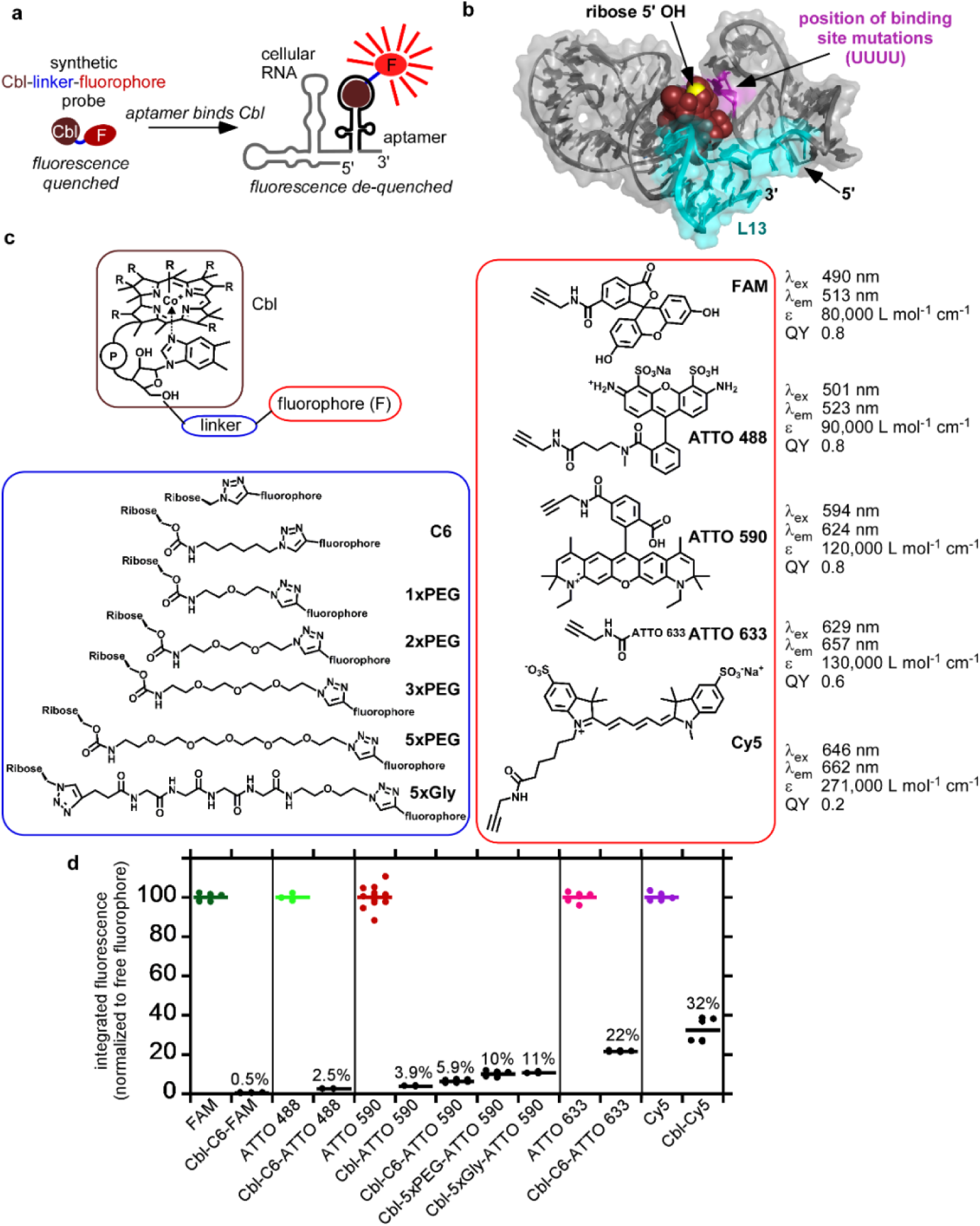
Covalent attachment of different fluorophores to Cobalamin (Cbl) results in fluorescence quenching, allowing for fluorescence turn-on of the probe upon binding to riboswitch RNA. (a) Principle of RNA-induced fluorescence turn-on for Cbl-fluorophore probes. Cbl (brown circle) acts as a quencher for the covalently attached fluorophore (red oval) due to proximity in space. Upon binding the RNA, Cbl is sterically separated from the fluorophore,resulting in de-quenching and fluorescence turn-on. (b) Structure of the Cbl riboswitch RNA (variant A, see Figure S4) bound to Cbl^30^. Loop P13 (teal) is at the 3’-end of the riboswitch. Cbl is shown in red spheres and the 5’-hydroxyl residues at the ribose moiety is shown in yellow.The four bases that were mutated to UUUU to abolish binding to Cbl are shown in magenta. (c) Synthetic Cbl-fluorophore probes used in this study. The organic linker was attached at the 5’-hydroxyl of the ribose and conjugated to alkyne variants of commercially available fluorophores via click chemistry, resulting in the triazole linkage between the linker and fluorophore. Note that the structure of ATTO 633 is proprietary. (d) Comparison of the fluorescence intensity of fluorophore *vs*. Cbl-fluorophore probes. Fluorescence spectra of 5 μM free fluorophores or Cbl-fluorophore probes were collected, the integrated intensity for each free fluorophore was set to 100% and the value for the Cbl-fluorophore probe was normalized relative to that of the corresponding fluorophore. Data are presented as mean for at least n = 3 independent measurements.

To assess how different fluorophores affect quenching, a series of probes with different fluorophores conjugated to Cbl via a 6-carbon chain (C6) was synthesized (Supplementary Note, Supplementary Fig. 2). The fluorescence of each probe was measured and compared to that of the free fluorophore (Fig. 1d, Supplementary Fig. 3). Cbl-C6-FAM and Cbl-C6-ATTO 488 retained only 0.5% and 2.5% fluorescence respectively, indicating that quenching, defined as reduction of the fluorescence signal, was highly efficient for probes with fluorophores in the green wavelength regime (λ_em_ ~520 nm). In contrast, probes with emission in the far-red range (λ_em_ ~660 nm) retained around 25% fluorescence (22% for Cbl-C6-ATTO 633 and 32% fluorescence for Cbl-Cy5), corresponding to weaker quenching (Fig. 1d). Cbl-C6-ATTO 590 emits in the red range (λ_em_ ~624 nm) and resulted in moderate quenching (5.9% residual fluorescence). Together, the results revealed a correlation between quenching efficiency and the excitation/emission wavelengths of the fluorophore, where quenching was the least efficient in the far-red and most efficient in the green wavelength range.

We systematically assessed how the length of the chemical linker affects quenching and found that increasing the linker length reduces quenching efficiency, consistent with similar observations in the literature^32^. Addition of a C6 linker (~10.5 Å, Supplementary Table 3) between Cbl and ATTO 590 resulted in higher residual fluorescence (5.9% for Cbl-C6-ATTO 590 vs. 3.9% for Cbl-ATTO 590) (Fig. 1d). In line with this trend, increasing the linker length further to five polyethylene glycol (PEG) units (~17.5 Å, Supplementary Table 3) or a 5x glycine linker (~21.4 Å, Supplementary Table 3) increased the residual fluorescence to 10% and 11%, respectively. Similar trends were observed when changing the linker length for probes with FAM as the fluorophore (Supplementary Fig. 3), confirming that quenching is most efficient when the fluorophore is close to Cbl.

### Variation of RNA properties modulate fluorescence turn-on

We hypothesized that binding of the RNA tag to the Cbl moiety of our probes would sterically separate fluorophore and quencher and thereby reduce quenching, resulting in an increase in fluorescence (Fig. 1a). To test this, the fluorescence signal of several probes in the presence and absence of a minimal truncated RNA tag (A_T_) was compared. We also included a control RNA that harbors four point mutations to abolish binding to Cbl (A_T,MUT_) (Supplementary Fig. 4). As expected, fluorescence increase was observed for all probes upon binding to A_T_ (Fig. 2a, b, Supplementary Fig. 5, see Supplementary Table 4 for a summary of fold turn-on results). Importantly, signal was not significantly changed in the presence of the negative control RNA A_T,MUT_ (Fig. 2a, b), and RNA A_T_ did not affect the fluorescence signal of the free fluorophore (Supplementary Fig. 6). Together, these observations indicate that fluorescence turn-on is specifically induced when RNA A_T_ binds the Cbl portion of the probe.

**Figure 2:**
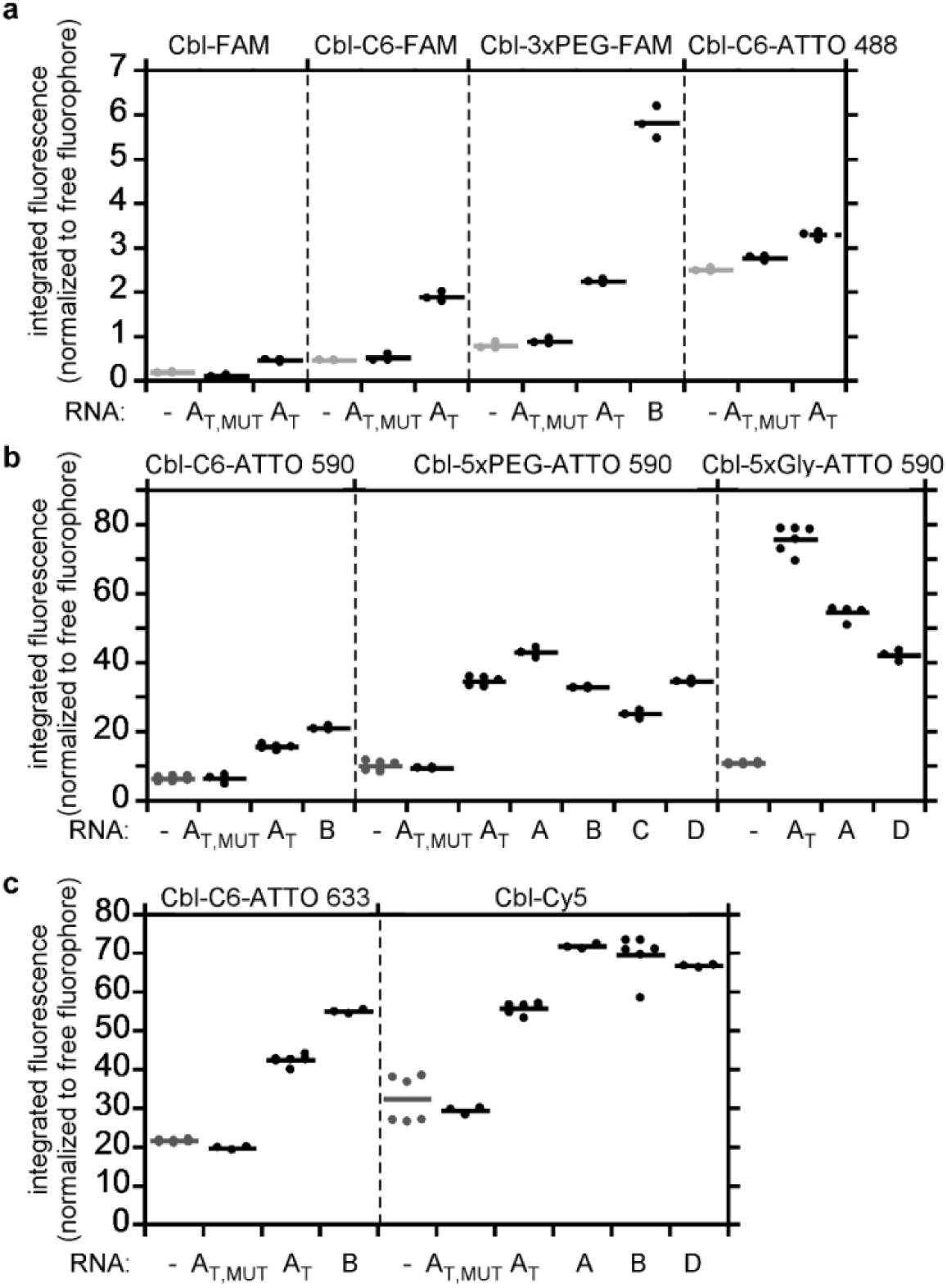
Cbl riboswitch RNAs induce fluorescence turn-on in Cbl-fluorophore probes *in vitro*. The fluorescence intensity of Cbl-fluorophore probes in the presence or absence of different RNAs was quantified as in Figure 1d and normalized relative to the intensity of the free fluorophore. The RNAs A, B, C and D are variants of Cbl-binding riboswitch sequences, where AT refers to a truncated version of A with linker region J1/3 and stem-loop P/L13 deleted (see also Figure 1b). The subscript MUT in A_T,MUT_ refers to four point mutations in the Cbl-binding site (see Figure 1b for the position of these residues). (a) Probes with fluorescence in the green wavelength range, (b) probes with fluorescence in the red wavelength range, (c) probes with fluorescence in the far red wavelength range. Data are presented as mean for n = 3 independent measurements (see Supplementary Table 4 for a summary).

We predicted that increasing the steric bulk of the RNA would promote greater separation between Cbl and fluorophore upon binding, leading to a larger fluorescence turn-on. To test this hypothesis, we compared fluorescence turn-on for Cbl-5xPEG-ATTO 590 in the presence of A_T_ *vs*. the full-length A that contains an additional structural element to increase bulkiness (Supplementary Fig. 4). Binding of Cbl-5xPEG-ATTO 590 to A compared to A_T_ resulted in a modest increase in fluorescence (35% *vs*. 43%, Fig. 2b, see also Supplementary Table 4 for a summary of fluorescence turn-on results). To further test the influence of RNA structure, we synthesized three additional riboswitch variants (B, C and D, Supplementary Fig.4) derived from other members of the Cbl riboswitch family. All three variants B, C and D include bulky features that could potentially introduce steric constraints when binding the probe (Supplementary Fig. 4). Variants B, C and D resulted in similar fluorescence turn-on compared with variant A for most probes tested (Fig. 2a, b). Together, bulky features appended to the aptamer only modestly affected the extent of fluorescence turn-on (Supplementary Table 4).

The observation that spectral properties of the fluorophore impacted the extent of quenching and fluorescence turn-on upon riboswitch binding suggests that Förster resonance energy transfer (FRET) may contribute to quenching, a hypothesis we evaluated by calculating the Förster radius and estimating the distance between the fluorophore and quencher for each of the probes (Supplementary Tables 2, 3, 5). The estimated distance between quencher and fluorophore for probes in the green wavelength range (FAM and ATTO 488 fluorophores) is significantly below the calculated values for R_0_, consistent with efficient quenching (< 3% residual fluorescence, Fig. 1d, 2a). In line with this model, even the bulkiest aptamers tested induced fluorescence turn-on that resulted in ~5% or less fluorescence compared with the free fluorophore (Fig. 2a), presumably because the length of the linker does not allow for separation of fluorophore and quencher beyond the R_0_ distance. While our theoretical estimates are consistent with a model where FRET contributes to quenching, it is noteworthy that conjugates in the far-red wavelength regime (λ_em_ ~660 nm) lack spectral overlap with the Cbl absorbance spectrum (Supplementary Fig. 7), yet moderate fluorescence quenching was still observed, suggesting that non-FRET mechanisms such as contact quenching or electron transfer must also contribute to fluorescence quenching. Probes with the ATTO 590 fluorophore have a Förster distance (R_0_ = 20 Å) close to the estimated distance between corrin ring and fluorophore (Supplementary Table 5), suggesting that ATTO 590 probes are particularly susceptible to large changes in fluorescence intensity for small distance changes.

### Biophysical characterization of RNA / probe complexes

We next identified RNA and Cbl-fluorophore probes with ideal photophysical behavior for cellular imaging and characterized their biophysical properties. We reasoned that shorter RNA tags would be less disruptive in RNA fusions. Therefore, we chose RNAs A and A_T_ for their small size (103 and 81 nucleotides, respectively) and induction of strong fluorescence turn-on *in vitro* (for example, 4.9x turn-on for A with Cbl-5xPEG-ATTO 590 and 7.3x turn-on for A_T_ with Cbl-5xGly-ATTO 590, Fig. 2, Supplementary Table 4). We also included aptamer D (130 nucleotides) for further *in vitro* characterization, since preliminary studies indicated high binding affinity to Cbl. Indeed, RNA tags A and D bind Cbl-5x-PEG-ATTO 590 tightly with a dissociation constant (K_D_) of 34 nM and 3 nM, respectively, comparable to the K_D_ for Cbl alone (Supplementary Fig. 8, Supplementary Table 6). The truncated riboswitch, A_T_, bound Cbl-5xPEG-ATTO590 with a lower affinity with a K_D_ of 1.3 μM.

We chose Cbl-fluorophore probes with red and far red fluorescent properties for further characterization for the following reasons. Red and far-red probes are desirable for live-cell imaging due to decreased cellular autofluorescence in this wavelength regime. While probes in the green wavelength range (for example Cbl-3xPEG-FAM, Fig. 2a) display efficient quenching and reasonable fluorescence turn-on (up to 7.4x fluorescence turn-on for Cbl-3xPEG-ATTO 488 with aptamer B, Fig. 2, Supplementary Table 4), the fluorescence signal after turn-on was low (only ~5% of fluorescence of free ATTO 488 in the presence of the best aptamer), raising concerns about the signal and contrast that would be achievable in live cell microscopy applications. Since FRET is likely a contributing factor to quenching in Cbl-ATTO 488 probes, higher fold turn-on will require a linker longer than the Förster radius of 35 Å (Supplementary Table 2), indicating the linker would need to be much longer than the longest linker in our study (5xPEG, 21.4 Å, Supplementary Table 3). Therefore, we chose red and far red fluorescent probes (Cbl-5xPEG-ATTO 590, Cbl-5xGly-ATTO 590 and Cbl-Cy5) since upon RNA binding, they exhibited robust turn-on and fluorescence closer to the free dye (Fig. 2b, c).

We then determined the quantum yield of probes Cbl-5xPEG-ATTO590, Cbl-5xGly-ATTO 590 and Cbl-Cy5 in the presence and absence of A and D (Supplementary Fig. 9, Supplementary Table 7). We observed a decrease in quantum yield for Cbl-fluorophore probes *vs*. free fluorophores and an increase in the presence of RNA, consistent with results from our plate reader fluorescence assay (Fig. 2b, c). In line with the changes in quantum yield, the fluorescence lifetime was reduced for Cbl-fluorophore probes *vs*. their free fluorophore counterparts and fluorescence lifetime increased in the presence of the RNA (Supplementary Fig. 10, Supplementary Table 8). Lastly, we assessed photostability of the red Cbl-5xPEG-ATTO 590 and the far red Cbl-Cy5 bound to RNA A *in vitro* under constant illumination. As seen by others^27^, the most widely used dye-binding RNA aptamer platform, DFHBI-1T-Broccoli, photobleaches to <20% fluorescence within a second. In contrast, Cbl-Cy5 bound to A retained >80% fluorescence even after 50 s constant illumination (>60% fluorescence after 50 s for Cbl-5xPEG-ATTO 590 bound to A) (Supplementary Fig. 11). Together, tight binding of our probes to the riboswitch variants A and D, robust increase in quantum yield upon RNA binding, as well as substantially slower photobleaching compared with the dye-binding aptamer Broccoli indicate favorable properties for cellular imaging applications.

### Visualization of mRNA dynamics in live mammalian cells

Our riboswitch-based tagging system includes several features that suggest broad applicability in live mammalian cell applications, prompting us to test our imaging platform by visualizing recruitment of β-actin mRNA (*ACTB*) to SGs in U2-OS cells tagged with the riboswitch variants (Fig.3, Supplementary Fig. 12). *In vitro* evolved dye-binding aptamers contain a G-quadruplex fold^37,38^ which has been shown to complicate RNA folding in mammalian cells^39^. Furthermore, ligands for evolved aptamers with the G-quadruplex fold may bind other RNAs with G-quadruplex folds^39^, have been shown to bind nucleic acids non-specifically^25^ and G-quadruplex structures may be disrupted by dedicated helicases^40^. Thus, to improve folding and stabilize against RNase degradation in cells, such aptamers often include a folding scaffold when fused to mammalian RNAs^25–27,41^. Two possible concerns of the folding scaffold are that it further increases the size of the RNA tag, and certain scaffolds increase undesired RNA processing in mammalian cells^42^. Our RNA tag did not require the tRNA folding scaffold^43^ and exclusion of the scaffold prevented unwanted processing (Supplementary Fig. 13) and reduced the overall tag size, prompting us to omit this scaffold for all further live cell experiments. The residual unquenched fluorescence of the three Cbl-fluorophore probes used in cellular studies (Cbl-5xPEG-ATTO 590, Cbl-5xGly-ATTO 590, Cbl-Cy5) revealed that in the absence of the riboswitch RNA, bead loading the probes in U2-OS cells^44–46^ gave rise to diffuse cytosolic and nuclear localization (Supplementary Fig. 14a-c). Treatment of U2-OS cells with arsenite induces formation of SGs that contain the marker protein G3BP1^47–49^, which can be tagged with GFP or the Halo-tag and subsequently labeled with red or far red fluorophores JF585, SiR594 or JF646^50^ (Supplementary Fig. 15). We verified that ACTB mRNA^47,48^ tagged with riboswitch variant A localized to G3BP1-positive SGs via fluorescence in situ hybridization (FISH) in fixed cells (Supplementary Fig. 16a), similar to endogenous ACTB mRNA (Supplementary Fig. 17). Together, tagging ACTB mRNA and visualizing recruitment to SGs is a robust system to assess our imaging platform in live mammalian cells.

Recruitment of ACTB mRNA tagged with the riboswitch RNA and colocalization with the fluorescently labeled marker protein G3BP1 was used to quantitatively validate the RNA imaging platform in live cells. To identify cells that produce the ACTB mRNA tagged with riboswitch variant A after transient transfection, we used a plasmid encoding for a blue nuclear NLS-TagBFP marker as a cotransfection marker and found that >90% of cells positive for NLS-TagBFP also produce the cotransfected mRNA (in this case mNeonGreen fused with RNA tag A, Supplementary Fig. 18). To visualize ACTB mRNA, cells were transfected with a plasmid to produce the A-tagged ACTB mRNA and the Cbl-Cy5 probe was loaded in live cells before inducing SGs by arsenite (Fig. 3). We observed robust accumulation of Cbl-Cy5 fluorescence in SGs upon arsenite treatment when 4 copies of A were used, but not when only one copy was used (Fig. 3b, d, Supplementary Fig. 20). When Cbl-Cy5 fluorescence was monitored over time upon arsenite treatment in cells containing ACTB-(A)4x mRNA, we detected formation of ACTB-(A)4x mRNA-containing SGs and monitored dynamics of SGs for ~ 50 min (Supplementary Fig. 19). Together, the SG visualization experiments demonstrate that ACTB mRNA recruitment to SGs and mRNA dynamics over time can be visualized via the riboswitch tag.

**Figure 3:**
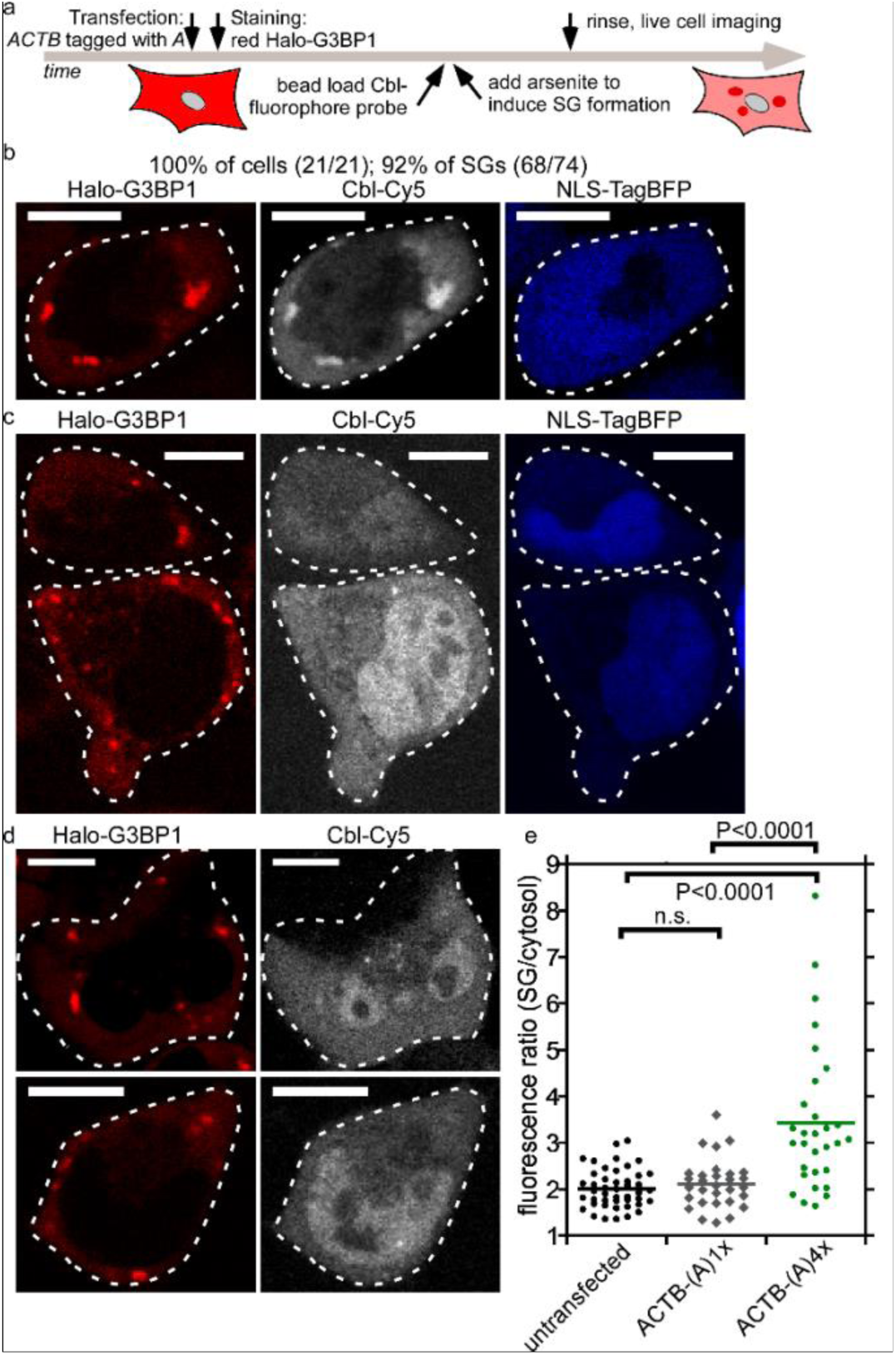
Monitoring ACTB mRNA localization to stress granules (SG) via Cbl-fluorophore probe binding to the RNA tag A. (a) Experimental strategy of labeling the Halo-tagged G3BP1 SG marker protein with the red fluorescent JF585 dye, followed by bead loading the Cbl-fluorophore probe, induction of SGs by arsenite for 30-45 min and live cell imaging. (b) U2-OS cells producing Halo-G3BP1 were transfected with ACTB-(A)4x and the transfection marker TagBFP. 24 h post transfection, cells were stained with the JF585 Halo dye. Shown are examples of representative live cells to assess colocalization of the SG marker protein Halo-G3BP1 and ACTB mRNA (3 experiments, 16 cells, 74 SGs). At least one SG was visible in all 16 cells (100%) and 92% of SGs were detectable. (c) The same experiment as in (b) was performed, except that ACTB-(A)1x was transfected (3 experiments, 14 cells, 30 SGs). Two representative live cells are presented. In 43% of the cells at least one SG was detectable and 40% of all SGs were detected in the Cbl-Cy5 channel. (d) The same experiment as in (b) was performed, except that ACTB-(A)4x was not transfected (2 experiments, 39 cells, 100 SGs).Two representative examples of live cells are presented. In 38% of the cells at least one SG was detectable and 32% of all SGs were visible in the Cbl-Cy5 channel. (e) Quantification of fluorescence signal accumulation in representative SGs. Scale bar = 10 μm. One way ANOVA (95% confidence limit), post hoc test (Tuskey HSD).

Importantly, the Cbl-Cy5 fluorescence signal remained diffuse throughout the cytosol in untransfected cells even when G3BP1-labeled SGs were induced (Fig. 3c, d), indicating that Cbl-Cy5 specifically binds to the A aptamer. In the absence of arsenite stress, the Halo-G3BP1 fluorescence signal was diffusely localized throughout the cytosol, and a smaller fraction of cells showed SG formation, in which case Cbl-Cy5 fluorescence localized to SGs (Supplementary Fig. 21). In this case, the SGs result from the process of transfection, which has been shown to induce SGs occasionally^51^. We also visualized ACTB mRNA tagged with 4 copies of the aptamer A and used the probe Cbl-5xGly-ATTO 590 and observed similar results (Supplementary Fig. 22, Fig. 4d). It is important to note that expression level of mRNA fusions and probe uptake efficiency (Supplementary Fig. 14a) are heterogeneous processes, which may explain the broad distribution of Cbl-Cy5 fluorescence increase in SGs (Fig. 3d). To directly confirm that Cbl-Cy5 fluorescence accumulation in SGs is due to the presence of tagged ACTB mRNA in SGs, we measured the formation of SGs in live cells via colocalization with the GFP-tagged G3BP1 marker protein and subsequently fixed cells and confirmed RNA tag colocalization with SGs by FISH (Supplementary Fig. 23). In summary, we demonstrated recruitment of ACTB mRNA to SGs in live cells using our riboswitch tagging system, enabling visualization of 100% cells containing at least 1 SG and 92% of all SGs, with the additional benefit of allowing visualization with two orthogonal colors (ATTO 590 and Cy5).

**Figure 4.**
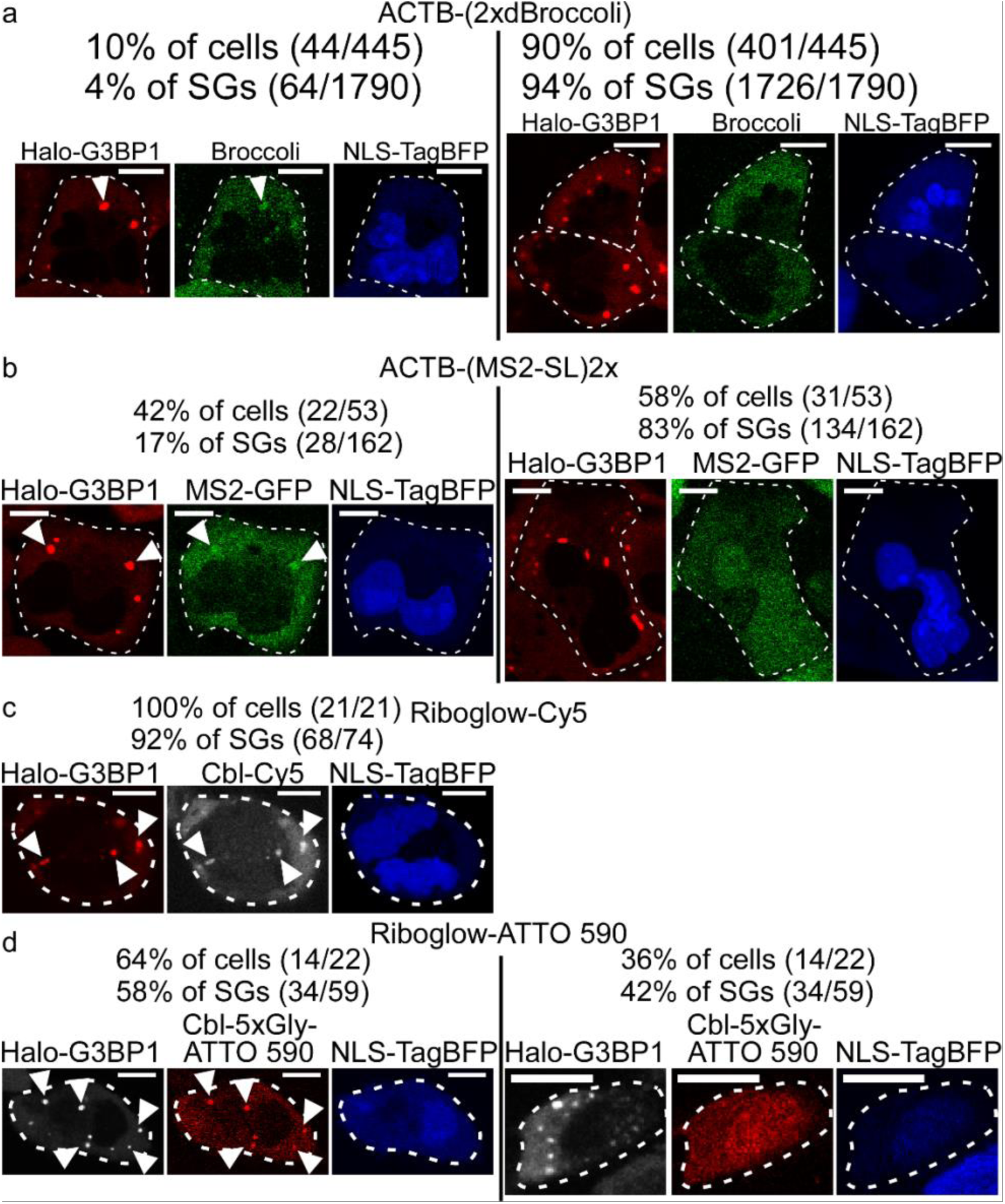
Comparison of ACTB mRNA imaging in stress granules (SG) by RNA tagging systems with four fluorophores per RNA. A plasmid encoding for tagged ACTB was transfected in U2-OS Halo-G3BP1 cells and with the NLS-TagBFP transfection marker. Halo-G3BP1 was labeled with a Halo dye for SG identification and SGs were induced by arsenite. (a) ACTB-(2xdBroccoli) was transfected, the Broccoli probe DFHBI-1T was added and cells were assessed for SGs in the green channel (445 cells, 3 experiments). In 10% of all transfected cells, at least one SG was detected and 4% of all SGs were detected (1790 SGs in 445 cells). (b) ACTB fused with two MS2 SL repeats was transfected with the transfection marker in U2-OS Halo-G3BP1 cells that stably produce MS2-GFP (in 42% of all transfected cells at least one SG was detected and 17% of all SGs were detected; 162 SGs total in 53 cells). (c, d) ACTB tagged with four copies of the riboswitch tag A was transfected with the transfection marker in U2-OS Halo-G3BP1 cells. (c) Cbl-Cy5 was loaded in cells (in 100% of all transfected cells at least one SG was detected in the Cy5 channel and 92% of all SGs were detected; 74 SGs total in 21 cells). (d) Cbl-ATTO 590 was loaded in cells (in 64% of all transfected cells at least one SG was detected in the Cy5 channel and 58% of all SGs were detected; 59 SGs total in 22 cells). Scale bar = 10 μm.

### Assessing performance of riboswitch tag versus existing RNA tags for live imaging

We next assessed performance of our riboswitch-based RNA tag for RNA imaging in live cells *vs*. commonly used RNA imaging platforms, namely the dye binding aptamer Broccoli^22,26^ and the MS2 system^9,10^. We used recruitment of tagged ACTB mRNA to SGs as a model system for this comparison as it represents a robust and well-established mRNA localization phenotype^47,48^. ACTB mRNA was tagged with the most robust Broccoli tag developed to date where a dimer of Broccoli dimers is integrated in the F30 folding scaffold^26^, such that each tag binds four fluorogens (called 2xdBroccoli, Supplementary Fig. 12, Table 1). A series of mRNA tags with the MS2 stem-loop repeats were generated, where 1, 2, 4 or 24 copies of the hairpin stem-loop were added (Supplementary Fig. 12, Table 1). It is important to note that dimers of MS2-GFP bind to each MS2 stem-loop^52^, such that mRNA with two copies of the MS2 stem-loop recruits a total of four GFP molecules per mRNA, leading us to choose ACTB-(MS2-SL)2x as a fair comparison with our platform where four copies of the RNA tag were used (Table 1). Tagging of ACTB mRNA with 2xdBroccoli or with up to 24 copies of the MS2 stem-loop does not affect mRNA localization to SGs (Supplementary Fig. 16b, c), leading us to use these constructs to test live mRNA recruitment to SGs.

We began by assessing ACTB mRNA recruitment to SGs using the 2xdBroccoli tag (Fig. 4a). We first confirmed that we could visualize Broccoli RNA when tagging 5*S* and U6 RNA using recommended experimental procedures^26^ (Supplementary Fig. 24). We then transfected a plasmid encoding for 2xdBroccoli-tagged ACTB mRNA and the NLS-TagBFP transfection marker in U2-OS cells that chromosomally produce Halo-G3BP1, induced SGs 24 h post transfection and added the DFHBI-1T probe to assess visualization of Broccoli-labeled SGs in the green fluorescence channel. In 90% of cells with the blue transfection marker that have Halo-G3BP1 labeled SGs in the red fluorescent channel, no corresponding SGs in the green Broccoli channel were detectable (401/445 cells, Fig. 4, Supplementary Fig. 25, right panel). In the remaining 10% of cells, green puncta corresponding to SGs (judged by colocalization with Halo-G3BP1 granules) were detectable (Supplementary Fig. 25, left panel), but only a minor fraction of all SGs were detected (4% of SG, 64/1790 SGs total). As seen for tagging with the riboswitch, transient transfection of the 2xdBroccoli-tagged ACTB mRNA in U2-OS cells alone occasionally induced G3BP1-positive SGs even in the absence of arsenite treatment (Supplementary Fig. 26). While rapid photobleaching of Broccoli is a well-established complication of the Spinach and Broccoli system^27,53^ (Supplementary Fig. 11), it is important to note that we used laser scanning confocal microscopy, a modality that resembles pulsed illumination shown to be ideal for Spinach and Broccoli imaging^53^. Nonetheless, we were unable to robustly visualize recruitment of ACTB mRNA tagged with a total of four Broccoli copies to SGs.

We then assessed ACTB mRNA recruitment to SGs via tagging with one, two, four or 24 copies of the MS2 stem-loop (Supplementary Fig. 27, Fig. 4b). A plasmid producing NLS-TagBFP was used as a transfection marker and SGs were induced 24 h post transfection for assessment of mRNA localization to Halo-G3BP1 labeled SGs. These experiments were performed in a variant of the U2-OS Halo-G3BP1 where NLS-MS2-GFP was also stably produced from the chromosome such that MS2-GFP remains nuclear in the absence of mRNA fused to MS2 stem-loops (Supplementary Fig. 28). As expected, localization of mRNA tagged with 24 copies of the MS2 stem loop to SGs was readily visualized via MS2-GFP recruitment to SGs in a large majority of cells that contained Halo-G3BP1 labeled SGs and that were also positive for the NLS-TagBFP transfection marker (86%, 43/50 cells, Supplementary Fig. 27d). When one, two or four copies of the MS2 stem-loop were used, visualization of ACTB mRNA localization to SGs was less efficient (28% of cells, 26% of all SGs for 1x MS2 SL tag; 42% of cells, 17% of all SGs for 2x MS2 SL tag; 45% of cells, 28% of all SGs for 4x MS2 SL tag, Fig. 4b, Supplementary Fig. 27). The fluorescence contrast for SGs over cytosolic background signal was markedly reduced when one, two or four copies of the MS2 stem-loop were used *vs*. 24 copies of the MS2 stem-loop, Supplementary Fig. 27). In summary, our riboswitch tagging system performed similarly to the 24x MS2 system with respect to reliably detecting localization of a candidate mRNA (ACTB) to SGs, while our platform outperforms both the Broccoli and MS2 system on a fluorophore by fluorophore basis (where four fluorophores per RNA are used, Fig. 4).

### Visualizing sequestration of non-coding U1 snRNA in U-bodies in live cells

The small size of our truncated RNA tag (81 nt for one copy of *A_T_*) opens up the possibility for tagging of ncRNAs such as snRNA U1 in live cells. Proper processing of U snRNA was found to depend on its length with an overall size limit of 200-300 nucleotides (nt)^54^, limiting the size of any U1 fusion tag to ~100 nt. To keep the RNA tag size as short as possible, we chose to use the A_T_ variant (81 nt, Table 1) and introduced this tag near the 5’ end of the U1 coding sequence (Supplementary Fig. 12), a position that was previously shown to be compatible with short RNA tags^55^. U snRNAs localize to nuclear Cajal bodies^56^ and U1 tagged with A_T_ can also localize to the Cajal body marker protein Coilin in HeLa cells (Supplementary Fig. 29a, b). Treatment of HeLa cells with Thapsigargin induced sequestration of the endogenous U1 snRNA and A_T_-tagged U1 in cytosolic U-bodies that contained the marker proteins DEAD-Box Helicase 20 (DDX20) and survival motor neuron (SMN) (Supplementary Fig. 29c, 30). When loading Cbl-fluorophore probes in live HeLa cells, non-specific puncta formation was observed for Cbl-Cy5 and for Cbl-5xGly-ATTO 590, but not for Cbl-5xPEG-ATTO 590, even at elevated probe concentrations (Supplementary Fig. 14d-g), leading us to choose Cbl-5xPEG-ATTO 590 for live U1 snRNA visualization. After Thapsigargin-treatment of cells transiently transfected with the A_T_-tagged U1, 24±10% of cells contained cytosolic puncta resembling U-bodies, whereas such puncta were only observed in 6±3% of untransfected cells (Fig. 5b). These results are in line with reports in the literature, where ~25% of HeLa cells were reported to contain U-bodies upon Thapsigargin-treatment^7^. Similarly, no cytosolic puncta were observed in the absence of Thapsigargin-treatment (Supplementary Fig. 31). To confirm that the cytosolic puncta we observed constitute U1 snRNA containing U-bodies, we transiently transfected the GFP-tagged U-body marker protein SMN together with the A_T_-tagged U1 and observed colocalization (Fig. 5c, Supplementary Fig. 32). Together, we conclude that our riboswitch-based imaging system allows for live cell visualization of small non-coding RNAs such as the snRNA U1.

**Figure 5.**
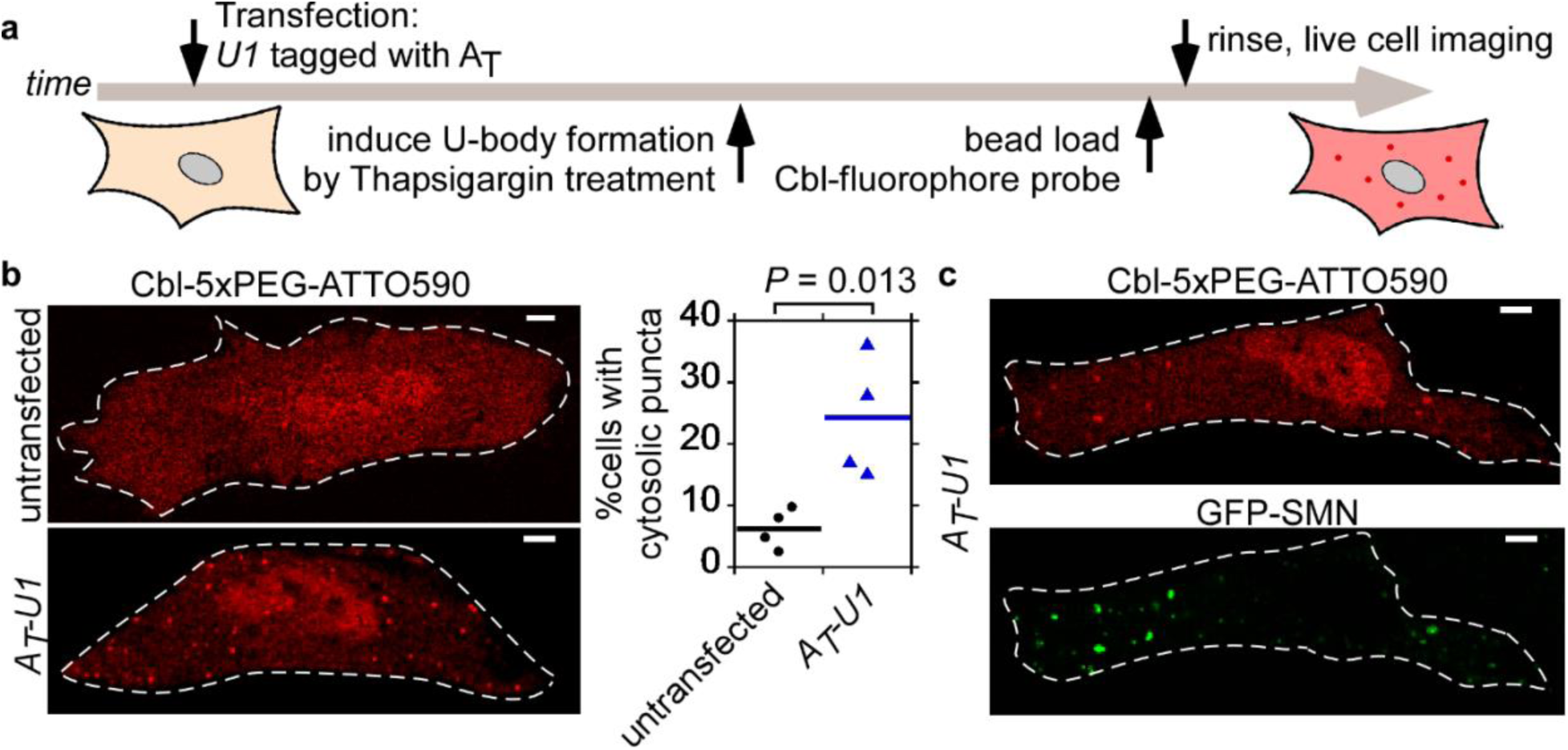
Monitoring cytosolic U-bodies via A_T_-tagged U1. (a) After transient transfection of A_T_-U1, U-bodies were induced by Thapsigargin treatment followed by live cell microscopy. (b) Cbl-5xPEG-ATTO 590 localization to cytosolic puncta in Thapsigargin-treated HeLa cells is more likely when A_T_-U1 was transfected (A_T_-U1: mean from 4 independent experiments / 326 cells; untransfected: mean from 4 independent experiments / 677 cells). One way ANOVA (95% confidence limit), post hoc test (Tuskey HSD). (c) Cytosolic puncta in Thapsigargin-treated cells expressing A_T_-U1 co-localize to GFP-SMN U-bodies. 3 independent experiments, 10 cells.Scale bar = 5 μm.

## Discussion

In this work we introduce a new riboswitch-based RNA imaging platform that we call ‘Riboglow’ characterized by several features that distinguish it from previous RNA-detection systems. First, the RNA tag is small and exploits the selective and high affinity (K_D_ in nM range) binding between a riboswitch and its cognate ligand, Cbl. Second, because the RNA tag binds the fluorescence quencher, Cbl, the system is compatible with a wide range of synthetic fluorophores spanning the green to far-red spectral range. The Riboglow-ATTO 590 and Riboglow-Cy5 probes used for live cell studies retain favorable photophysical properties, including slow photobleaching, of the parent dyes when bound to RNA. Indeed, we were able to visualize tagged mRNA over 50 min without detectable photobleaching (Supplementary Fig. 19). Third, our RNA tag is not subjected to undesired processing and hence does not require the tRNA-like scaffold used for other dye-binding aptamers. Fourth, cobalamin riboswitches include a large family of RNA sequences that all bind Cbl^57^ that can be exploited for future optimization and customization of the Riboglow platform. Fifth, the fluorescence lifetime of ATTO 590 and Cy5 in the context of our probes varies when the RNA binds (Supplementary Fig. 10), raising the possibility of using this system for fluorescence lifetime imaging. Finally and perhaps most importantly, we were able to visualize recruitment of RNA polymerase II-dependent transcripts (mRNA and snRNA) to RNP-granules, where the short size of the RNA tag (~80 nts for RNA tag A_T_) enabled tagging and visualization of U1 snRNA in live cells for the first time.

To compare the performance of Riboglow to existing RNA imaging platforms, we characterized recruitment of mRNA to SGs in live cells. Riboglow is most similar to dye-binding systems (Spinach^22^, Broccoli^23^ and Mango^24,25^) in terms of concept, size of the RNA tag, and molar mass of the probe. We used a two-fold repeat of dimer-Broccoli as our point of comparison because it is the brightest of the Spinach/Broccoli family of dye-binding aptamers. The Mango system, which was published recently, appears to exhibit properties similar to Broccoli and the dye is not commercially available, making it challenging to test. Compared to Riboglow (92% of SGs for Riboglow-Cy5, Fig. 3, 58% of SGs for Riboglow-ATTO 590, Supplementary Fig. 22), only a small minority of SGs were detected via the Broccoli-labeled mRNA (4% of SGs, Supplementary Fig. 23). This is perhaps not surprising because dye-binding aptamers have not yet been used to detect RNA polymerase-II transcripts, such as mRNA. Rather, these systems have been applied to RNA polymerase-III transcripts^25,27^ which are expressed at much higher levels, particularly in HEK 293 cells. It is noteworthy that our riboswitch tagging system displays much weaker fold fluorescence turn-on *in vitro* than the ~1,000-fold *in vitro* fluorescence enhancement for both Broccoli^23^ and Mango^24,25^ (Fig. 2). However, cellular contrast defined as fluorescence signal in RNA-granules *vs*. cytosolic background was comparable for our platform (~3-4-fold, Fig. 3d) and Mango-tagged 5*S*-RNA and U6-RNA foci (~2-3-fold fluorescence turn-on in fixed mammalian cells^25^). The *in vitro*-evolved dye-binding aptamers all feature G-quadruplex RNA folds^38,39^, and it has been shown that the dyes can bind other G-quadruplex RNAs non-specifically in cells^39^, possible contributing to diminished cellular contrast. It is important to note that other factors besides *in vitro* enhancement contribute to cellular contrast, including robust folding of the RNA tag in the cellular environment, temperature- and salt-dependent stability of the RNA-probe complex, and probe photobleaching properties. Indeed, Broccoli is less photostable than the Riboglow probes which could contribute to its poor performance in live cell studies. We observed poor performance of Broccoli, even though it is worth noting that pulsed illumination modalities such as the laser scanning confocal setup we used were shown to markedly improve microscopy image acquisition for Spinach^53^.

The MS2/PP7 system is the gold standard for detection of mRNA dynamics in mammalian cells^9–11^. Indeed, we found that 1x, 2x, 4x, and 24x copies of the stem-loop all permitted visualization of mRNA recruitment to SGs. Not surprisingly, the 24x tag yielded the strongest fluorescence contrast and led to detection of the greatest number of SGs (86% of cells with at least 1 SG were detected, 76% of all SGs were detected, Supplementary Fig. 27d). Our Riboglow platform outperformed the MS2 system in a fluorophore by fluorophore comparison where four GFP molecules were bound to each RNA and the Riboglow platform used four fluorophore-binding RNA tags (Table 1, Fig. 4). Riboglow even compared favorably with the 24x MS2 system, yielding robust detection of SGs in transfected cells (92% of SGs for Riboglow-Cy5, 58% of SGs for Riboglow-ATTO 590) and robust fluorescence contrast of labeled mRNA in SGs (on average 3-4-fold, Fig. 3d).

There are some properties of Riboglow that are less favorable when compared to other RNA-tagging systems. While the Broccoli probe DFHBI-1T and Mango probe TO1-Biotin are cell-permeable, we use bead loading for cellular Cbl-fluorophore probe uptake. Although we have found bead loading to be a robust and simple procedure that is routinely used in other live cell imaging applications^44,45^, this process adds an additional step to the imaging process. Utilization of the natural cobalamin uptake route may present an alternative in the future ^58,59^. Additionally, mammalian cells contain at least four known Cbl-binding proteins and it is possible that our Cbl-fluorophore probes could bind to these proteins, yielding fluorescence turn-on (much like dyes binding to G-quadruplexes in cells). Although we observe robust 3-4x contrast when tracking recruitment of RNA to RNA-protein granules, if non-specific binding is a problem for other applications, the Cbl probes could be further engineered to enhance specificity.

Overall, the Riboglow platform compares favorably in a direct side-by-side comparison with existing RNA imaging platforms and presents an orthogonal approach for live cell RNA imaging applications. The highly modular nature of the Riboglow platform currently allows for multicolor imaging and detection of mRNAs as well as small non-coding RNAs, while presenting a system that is ideally set up for straightforward future customization to include desired features for RNA imaging.

## Acknowledgements

The authors would like to acknowledge financial support from the Human Frontiers Science Project and NIH Director’s Pioneer Award GM114863 (to AEP). We acknowledge support from the National Science Centre, SYMFONIA DEC-2014/12/W/ST5/00589 to Dorota Gryko and Aleksandra J. Wierzba and from the National Institutes of Health (5R01 GM073850) to Robert T. Batey. We thank Luke Lavis for providing fluorophores JF585, SiR594 and JF646 for Halo staining, Siddharth Shukla and Jennifer Garcia for helpful discussions and for providing cell lines, antibodies and plasmids, Denise Muhlrad, Jason Lee and Maria Lo for technical expertise, and to Jens Eberhard and Jeremiah Gassensmith for helpful discussions. The imaging work was performed at the BioFrontiers Institute Advanced Light Microscopy Core, whose Nikon A1R microscope was acquired by the generous support of the NIST-CU Cooperative Agreement award number 70NANB15H226.

## Author contributions

EB, JTP, RTB and AEP conceptualized and designed the study. JTP and RTB rationally designed riboswitch variants. EB, JTP, RTB, AJW, DG and AEP designed organic probes. AJW and MC synthesized organic probes. JTP and ZH purified riboswitch variants for *in vitro* work. EB performed *in vitro* work and designed and performed cellular work and analyzed data with input from all authors. DB constructed plasmids and assisted with cellular work. STH performed *in vitro* fluorescence lifetime and bleaching experiments. JRW made the Halo-G3BP1 U2-OS cell line. RP and RJ provided critical advice. EB and AP wrote the manuscript with edits from all authors.

## Online Methods

### Synthesis of fluorophore probes

Descriptions of synthesis and characterization of probes can be found in the Supplementary Note. All probes are derivatives of CN-Cbl and structures of all probes used in this study are shown in Supplementary Figure 1. Supplementary Table 1 provides a summary of their photophysical properties.

### RNA synthesis and preparation

For all *in vitro* experiments, DNA templates were amplified using recursive PCR and transcribed by T7 RNA polymerase using established methods.^60^ Transcription reactions were purified using the appropriate percentage denaturing polyacrylamide gel (8 M urea, 29:1 acrylamide:bisacrylamide) based on RNA length. Transcripts were visualized by UV shadowing, excised from the gel and soaked overnight in 0.5x TE buffer (10 mM Tris-HCl, pH 8.0, 1 mM EDTA) at 4°C. RNAs were buffer exchanged into 0.5x TE and concentrated using centrifugal concentrators (Amicon) with the appropriate molecular weight cutoff. RNA concentration was determined using the absorbance at 260 nm and molar extinction coefficients calculated as the summation of the individual bases. Sequences and secondary structures of all riboswitch RNAs used in this study are shown in Supplementary Fig. 1 and the published sequence for the Broccoli sequence was used (5’-GGA GCG CGG AGA CGG TCG GGT CCA GAT ATT CGT ATC TGT CGA GTA GAG TGT GGG CTC CGC GC)^26^.

### Absorbance measurements and quantum yield determination

Absorbance spectra were collected using a Cary 500 UV-VIS-NIR spectrophotometer and buffer subtracted and measurements were reproducible compared with literature data. The quantum yield (Q) of probes was determined by comparison with the published quantum yield of ATTO 590 (0.80, Atto tec) or the published quantum yield of Cy5 (0.28, Lumiprobe). All experiments were conducted in RNA buffer (100 mM KCl, 10 mM NaCl, 1 mM MgCl_2_, 50 mM HEPES, pH 8.0). First, absorbance was determined at the excitation wavelength (Supplementary Table 1) for a dilution series of free fluorophore and the indicated Cbl-fluorophore probes in the presence and absence of different RNAs. To ensure saturation of binding, the concentration was chosen such that the final concentration of RNA was above 5 µM in the most diluted sample. The fluorescence spectrum at the emission range indicated in Table S1 for each sample was then recorded using a PTI-fluorimeter (1 nm steps, 1 s integration time, 2 nm slits for the excitation and 4 nm slits for detector). After subtracting the buffer background signal, the fluorescence signal for each sample was integrated and plotted *vs*. the absorption.The steepness of the resulting linear plot for each dilution series reports on the quantum yield relative to the reference of the free fluorophore (Supplementary Table 7). These measurements were done once for the Q determination, and spectra were comparable with absorbance / fluorescence measurements done for fold fluorescence measurements.

### Fluorescence lifetime measurements

The fluorescence lifetime of Cbl-fluorophore probes in the presence and absence of RNA (see Supplementary Table 8) was measured in RNA buffer (100 mM KCl, 10 mM NaCl, 1 mM MgCl_2_, 50 mM HEPES, pH 8.0) using Time-Correlated Single Photon Counting (TCSPC). A PicoQuant FluoTime 100 fluorescence spectrometer and PicoQuant Picosecond Pulsed Diode Laser Heads with wavelengths 561 nm (LDH-D-TA-560) and 640 nm (LDH-P-C-640B) were used in this measurement. Data were iteratively reconvoluted from the Instrument Response Function with one to three exponential decay functions, and the intensity weighted average lifetime values are reported.

### Photobleaching measurements

For photobleaching experiments, indicated probes and RNA (Supplementary Table 9) were mixed before about 3 to 4x of microscope immersion oil (Olympus immersion oil type-F) was added and the solution was emulsified to generate droplets. Samples were prepared in RNA buffer for Cbl-fluorophore samples (100 mM KCl, 10 mM NaCl, 1 mM MgCl_2_, 50 mM HEPES, pH 8.0) and in Broccoli buffer (40 mM HEPES pH 7.4, 100 mM KCl, 1 mM MgCl 2) for the DFHBI-1T / Broccoli sample. The droplets were sandwiched between a glass slide and a coverslip. The sample was placed on the sample stage of a Nikon Ti-E HCA widefield fluorescence microscope with the coverslip facing a 20× objective lens. All samples were continuously illuminated throughout the experiment at the irradiance reported in Supplementary Table 9. We assessed photostability by fixing the excitation rate, i.e. the number of absorbed excitation photons were the same for the Cbl-fluorophore / RNA samples and the DFHBI-1T / Broccoli sample. The high irradiance for the DFHBI-1T / Broccoli sample is a result of its low extinction coefficient. Images were acquired every 50 ms for DFHBI-1T / Broccoli and every 250 ms for others for 3 seconds, then every second for 47 seconds or longer, and the exposure time of each image was 15 ms for DFHBI-1T / Broccoli and 40 ms for others. A total of 6 to 15 droplets were analyzed for each sample.

### *In vitro* fluorescence measurements

The concentration of Cbl-fluorophore probes, free fluorophores and free Cbl was determined using extinction coefficients listed in Supplementary Table 1. All *in vitro* experiments were conducted in RNA buffer (100 mM KCl, 10 mM NaCl, 1 mM MgCl_2_, 50 mM HEPES, pH 8.0). For the Cbl-fluorophore probes, the extinction coefficient of the fluorophores was used to determine the concentration. The shape of the absorption spectra did not change significantly for the conjugated probes compared with the sum of spectra for free fluorophores and free Cbl (Supplementary Fig. 2). The fluorescence intensity of each probe was measured in the presence or absence of RNA in RNA buffer as technical triplicates in a Tecan Safire-II fluorescence plate. The concentration of the probe was 0.25 μM or 0.5 μM and the RNA concentration was at least 5 μM, significantly above the dissociation constant (K_D_) of the RNAs tested^30,31^. Samples were incubated for at least 20 min at room temperature in the dark to allow for Cbl-binding of the RNA prior to data collection. The excitation wavelength for each probe is listed in Supplementary Table 1 and the emission spectrum was collected in 1 nm increments for the range listed in Supplementary Table 1. Each emission spectrum was buffer subtracted, integrated and normalized to the signal of the free fluorophore at the same concentration.Resulting % fluorescence measurements were reproducible for measurements on different days. Representative fluorescence spectra of probes in the presence and absence of RNA are shown in Supplementary Figure 5.

### Isothermal titration calorimetry (ITC)

ITC experiments were performed using protocols previously established and described^30,31^. Briefly, the RNA was dialyzed overnight at 4°C into RNA buffer (100 mM KCl, 10 mM NaCl, 1 mM MgCl_2_, 50 mM HEPES, pH 8.0) using 6-8000 Dalton molecular weight cutoff dialysis tubing (Spectra/Por). The dialysis buffer was used to dissolve Cbl and Cbl-5xPEG-ATTO590, and to dilute RNA to the desired concentration. The concentration of Cbl and Cbl-fluorophore probe was determined using the extinction coefficients listed in Supplementary Table 1. Titrations were performed at 25°C using a MicroCal ITC_200_ microcalorimeter (GE Healthcare) and data were fit using the Origin software suite as previously described^61^.

### Estimation of theoretical photophysical properties and distances in probes

The overlap integral J(λ) of each fluorophore with Cbl absorbance was calculated from equation (1)^62^:

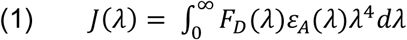

where F_D_ is the emission spectrum of the fluorophore normalized to unity and ε_A_ is the extinction coefficient of the acceptor (Cbl) in M^−1^ cm^−1^ using a MATLAB script (a|e - UV-Vis-IR Spectral Software 1.2, FluorTools,www.fluortools.com). To include the entire fluorescence emission spectrum of each fluorophore (Supplementary Fig. 6), fluorophores were excited 10 nm below the typical (Supplementary Table 1) excitation λ and emission was collected 10 nm above the excitation λ using a PTI-fluorimeter (1 nm steps, 1 s integration time, 2 nm slits for the excitation and 4 nm slits for detector). Cbl absorbance and fluorophore emission spectra (shown in Supplementary Fig. 7) and were converted in units of M^−1^ cm^−1^ (for absorbance of Cbl) and to unity (for emission of fluorophores) to calculate J(λ). The Förster distance R_0_ was calculated from equation (2)^62^

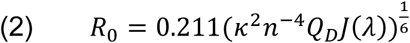

where κ2 is a factor describing the relative orientations of the transition dipoles and was assumed as κ2 = 2/362 and the refractive index n was assumed to be 1.4 for aqueous solutions. The quantum yield Q of each donor fluorophore is available from the manufactures of the fluorophores and is listed in Supplementary Table 2.

The maximal distance between quencher and fluorophore in each probe was estimated as the length of the chemical linker (listed in Supplementary Table 3) plus the distance between the 5’ hydroxyl moiety of Cbl (where the linker is attached) and the corrin ring in the structure of Cbl^30^, assuming that this region of Cbl harbors quenching properties. The distance between 5’ hydroxyl moiety and the corrin ring was estimated to be 9 Å from the Cbl crystal structure^30^.This distance was added to each linker lengths, resulting in the distances listed in Supplementary Table 5.

### Construction of plasmids for mammalian expression of RNA tag variants

Plasmids to produce RNA tags fused with RNA of interest in mammalian cells were constructed by standard molecular cloning techniques (see Supplementary Fig. 12 for an overview and Supplementary Fig. 1 for RNA sequences). To construct mRNA fusions, the commercially available vector *pmCherry-C1* (Clontech) was used as a starting point, since it harbors the CMV promoter for strong expression in mammalian cells. A *Nhe*I restriction site was introduced immediately following the stop codon by site directed mutagenesis. Different RNA sequences were inserted between the stop codon and the polyA site using existing *Kpn*I and *Bam*HI restrictions sites. Inserts encoding for RNA tags were purchased as g-blocks or ultramers depending on sequence length with the appropriate restriction site overhangs from IDT. Sequences for RNA tags are summarized in Supplementary Fig. 1 and the tRNA scaffold previously used in Spinach constructs^22^ was used. For ACTB mRNA fusions, the coding sequence of *mCherry* was then replaced with the *ACTB* coding sequence. The ACTB coding sequence was PCR amplified from a plasmid producing a mNeonGreen-actin fusion (sequence was verified to be identical with Homo sapiens beta actin (ACTB), NCBI Reference Sequence NM_001101.3) by standard restriction based cloning using *Nhe*I restriction sites flanking the coding sequence of the *mCherry* gene. As for mCherry fusions, the RNA tags were purchased as ultramers or gblocks from IDT and added by standard restriction based cloning using existing *Kpn*I and *BamH*I sites. The monomeric tags were introduced by ligating the first g-block into the plasmid with *Kpn*I / *BamH*I restriction sites resulting in ACTB-(A)1x. Additional copies were introduced sequentially, where the second and third copy was encoded on a g-block with *Kpn*I / *Kpn*I and *BamH*I / *BamH*I restriction sites, respectively, resulting in ACTB-(A)2x and ACTB-(A)3x. ACTB-(A)4x was the result of the third ligation step, where colonies were screened for insertion of more than one copy of A. ACTB-(A_T_)4x was constructed using the same strategy.

The 2xdBroccoli sequence^26^ to generate ACTB-2xdBroccoli was purchased as a g-block with *Kpn*I and *Bam*HI overhangs and ligated downstream of the ACTB sequence analogous to riboswitch RNA tags described above. Plasmids pAV5S-F30-2xdBroccoli and pAVU6+27-F30-2xdBroccoli^42^ were a gift from Samie Jaffrey (Addgene plasmid # 66845 and # 66842). The 1x and 2x copies of MS2 stem-loop (SL) sequences were purchased as g-blocks with *Kpn*I and *Bam*HI overhangs and ligated downstream of the ACTB sequence analogous to riboswitch RNA tags described above. The 4x MS2 SL sequence was generated by ligating another 2x MS2 SL g-block sequence with *Kpn*I restriction sites on both ends in the ACTB-(MS2-SL)2x plasmid.The 24xMS2 SL tag was built by first replacing the RNA tag downstream of *ACTB* with a short g-block sequence consisting of *Not*I and *Pme*I sites and flanking *KpnI* and *Bam*HI sites using *Kpn*I and *Bam*HI sites. Plasmid ACTB-(MS2-SL)24x was then produced by cutting the 24x MS2-SL repeat from plasmid PGK1-24×MS2-SL^12^ with *Not*I and *Pme*I sites and ligating it downstream of the ACTB sequence via *Not*I and *Pme*I sites.

The existing plasmid pU1(human)^63^ to produce the A_T_-tagged U1 snRNA was modified to add the sequence of the RNA tag A_T_ immediately following the first 11 nucleotides of U1, analogous to previous U1 snRNA fusions^55^. Briefly, the parent plasmid was digested with two existing unique restriction sites (*Bgl*II is upstream of the U1 coding sequence and *Pst*I is in the 3’ region of the U1 coding sequence). An insert that contains the sequence for A_T_ and the surrounding U1 coding sequence including both restriction sites was purchased as a g-block from IDT and ligated, resulting in A_T_-U1. All plasmids were verified by sequencing.

### Cell culture and cell lines

U2-OS cells (received from Professor Roy Parker^64^), HeLa cells and 293T cells (both purchased from ATCC) were maintained in Dulbecco’s modified eagle medium (DMEM, Gibco) supplemented with 10% fetal bovine serum (FBS; Gibco) at 37°C with 5% CO_2_. To generate a U2-OS cell line that stably produces GFP-G3BP1, the GFP-G3BP1 coding sequence was PCR amplified from a peGFP-C1-based cloning vector (received from Professor Roy Parker) using *Eco*RI and *Not*I restriction site overhangs. The PCR product was ligated into a Piggybac Dual Promoter plasmid that includes a Puromycin resistance cassette for selection and a CMV promoter for GFP-G3BP1 expression (System Biosciences, Catalog # PB510B-1), resulting in plasmid pGFP-G3BP1-Piggybac. This plasmid was sequence verified. U2-OS cells were chemically transfected with pGFP-G3BP1-Piggybac and Super PiggyBac Transposase expression vector (System Biosciences, Catalog # PB220PA-1) using the *Trans*IT transfection system according to manufacturer recommendations (Mirus). Selected for genomic integration using was started 3 days after transfection using 1 μg/mL Puromycin and continued for 7-10 days. After selection, U2-OS cells stably producing GFP-G3BP1 were FACS enriched for the brightest 30% of GFP-fluorescent cells. After FACS enrichment, cell aliquots were used for up to 10 additional passages.

To generate a U2-OS cell line that stably produces MS2-GFP, plasmid pMS2-GFP (a gift from Robert Singer, Addgene plasmid # 27121) was PCR amplified with *Xba*I and *Not*I restriction site overhangs and ligated into the Piggybac Dual Promoter plasmid described above. Transfection in U2-OS cells with a Halo-G3BP1 genetic background, selection and FACS enrichment was done as described above.

Introduction of an N-terminal 3xFLAG-HALO epitope tag to the endogenous G3BP1 protein was performed as previously described^65^ with the following modifications (Supplementary Fig. 33). First, two single-strand breaks were generated flanking the translational start site of G3BP1 with dual CRISPR-Cas9 nickases (D10A) in U2-OS cells. Correct targeting of crispr guides was confirmed by T7 Endonuclease assay (NEB). Cells were selected with Puromyocin to enrich for edited cells containing the sequence for the tags and the LoxP-flanked, SV40-driven Puromycin expression cassette. Removal of the Puromyocin expression cassette and in frame expression of 3xFlag-HALO was achieved by transient transfection of eGFP-Cre plasmid. Transiently transfected cells were enriched by FACS. Correctly edited cells were confirmed by PCR, Western blot and immunofluorescence using standard protocols.

### Halo-staining

Halo dyes JF585, SiR594 and JF646 were gifts from Professor Luke Lavis. Cells were incubated with 1 μM Halo dye in DMEM/5%FBS media for 20 min at 37°C / 5% CO_2_. Unbound dye was washed out by replacing the media first with PBS and then with DMEM/5%FBS media before further treatment was performed (arsenite treatment, bead loading, see below).

### Bead loading

Cells were seeded in home-made imaging dishes (35 mm diameter) with a ~10 mm center hole covered by cover glass (No. 1.5, VWR). To introduce the probe, the culture media was removed and the Cbl-fluorophore probe (3 μL of a 50 μM stock in PBS for U-body imaging in HeLa cells, 3 μL of a 0.5 μM stock in PBS for mRNA imaging via Cbl-Cy5 in U2-OS cells and 3 μL of a 5 μM stock in PBS for mRNA imaging via Cbl-5xGly-ATTO 590 in U2-OS cells) was added directly on the cell in the center of the imaging dish. Microbeads were sprinkled onto the cells^44–46^ and the dish was tapped on the cell culture surface 7-8 times. Standard culture media was added immediately. For stress granules imaging, media was supplemented with 0.5 mM sodium arsenite.

### Stress granule (SG) assay

U2-OS cells stably producing GFP-G3BP1 or genomically producing Halo-G3BP1 were seeded at 0.25×10^6^ cells in imaging dishes. One day after seeding, cells were chemically transfected with typically 2 μg plasmid DNA (1 μg of the *ACTB* plasmid fused with an RNA tag of interest mixed with 1 μg of the transfection marker pNLS-TagBFP (in pcDNA)) using the *Trans*IT transfection system following manufacturer recommendations (Mirus). On the next day, cells were stained and bead loaded with the Cbl-fluorophore probe as described above. The media that was added after bead loading contained 0.5 mM sodium arsenite and cells were incubated at 37°C / 5% CO_2_ for 30-45 min to induce stress granules. Cells were rinsed once in PBS and Fluorobrite media (Gibco), supplemented with 0.5 mM sodium arsenite, was added for live cell imaging. For correlative live / fixed imaging where cells were first imaged live, followed by fixation and FISH / immunofluorescence imaging of the same cells, the following modifications were made to the protocol above. Instead of home-made imaging dishes, gridded imaging dishes (MatTek) were used. Dishes were coated with 1 μg/mL fibronectin (Sigma) for 4 hours and fibronectin was rinsed once with full media before cells were seeded. Because plasmid ACTB-(AT)4x was used, a higher concentration of Cbl-5xPEG-ATTO 590 was bead loaded (3 μL of 50 μM probe).

### U-body assay

HeLa cells were seeded at 0.1 - 0.15 × 10^6^ cells per imaging dish. One day after seeding, cells were chemically transfected with 1 μg plasmid DNA (A_T_-U1) using the *Trans*IT transfection system following manufacturer recommendations (Mirus). For experiments with marker proteins, 0.25 μg of GFP-SMN^66^ (a gift from Greg Matera, Addgene plasmid #37057) was co-transfected. For colocalization experiments with Coilin, 0.5 μg pEGFP-coilin^67^ (a gift from Greg Matera, Addgene plasmid #36906) was transfected with or without co-transfection of 0.5 μg plasmid DNA encoding for A_T_-U1. Imaging or fixation was performed the following day.For live cell imaging, cells were treated with 10 μM Thapsigargin for 3 hours before bead loading with the Cbl-fluorophore probe and imaged within 1 hour after bead loading. For immunofluorescence or FISH analysis, cells were fixed 3-4 hours after Thapsigargin treatment.

### Fluorescence microscopy and image analysis

Fluorescence microscopy on live and fixed cells was performed on a Nikon A1R Laser Scanning Confocal Microscope with a 100x oil objective (1.45 NA, Plan Apo I), a pixel size of 0.25 μm and an integration time of 2.2 μsec unless otherwise noted below. Images were acquired at 16-bit depth with Nikon Elements Software and processed in ImageJ2 using the Fiji plugin. All live images were acquired with an environment chamber at 37°C. Laser lines used were 405 nm (for nuclear staining and NLS-TagBFP imaging), 488 nm (for GFP-tagged proteins), 561 nm (for ATTO 590 in Cbl-fluorophore probes and Alexa 546, Alexa 594 and Alexa 568 in FISH probes and secondary antibodies and for Halo-dyes JF595 and SiR594) and 638 nm (for Cy5 and Halo-dye JF646).

For live imaging of SGs tagged with the aptamer A bound to the Cbl-Cy5 probe in U2-OS cells, the pinhole was 67.7 μm and the laser settings were as follows. For the DAPI channel, the laser was set at 1.0 (HV = 100), for the TRITC channel the laser was set at 0.4 (HV = 20), for the Cy5 channel the laser was set at 12.0 (HV = 100). The signal was integrated 4x for each acquisition. For live imaging of SGs tagged with the aptamer A bound to the Cbl-5xGly-ATTO 590 probe in U2-OS cells, the pinhole was 67.7 μm and the laser settings were as follows. For the DAPI channel, the laser was set at 1.0 (HV = 100), for the TRITC channel the laser was set at 1.0 (HV = 80), for the Cy5 channel the laser was set at 5.0 (HV = 100).

For correlative imaging to visualize the same stress granules samples live and fixed, the pinhole size was 67.7 μm. In this case, the laser power for live imaging was 1.0 (HV = 40) for the 488 laser line and 7.0 (HV = 110) for the 638 nm laser line (the signal was not integrated). Fixed images for correlative imaging of stress granules were collected using a 40x Plan Apo Air objective (26.8 μm pinhole) with Nyquist sampling at 0.16 μm per pixel. The laser power for the 488 nm laser line was 2.0 (HV = 40) and the laser power was 1.0 for the 461 nm laser line (HV = 40).

For live imaging of SGs tagged with 2xdBroccoli in U2-OS cells, the pinhole was 67.7 μm and the laser settings were as follows. For the DAPI channel, the laser was set at 5.0 (HV = 110), for the GFP channel the laser was set at 1.0 (HV = 110), for the TRITC channel the laser was set at 1.0 (HV = 40). For live imaging of SGs tagged with MS2 stem-loop repeats in U2-OS cells, the pinhole was 67.7 μm and the laser settings were as follows. For the DAPI channel, the laser was set at 5.0 (HV = 110), for the GFP channel the laser was adjusted for optimal contrast without detector saturation (~1.0-2.5) (HV ~60-100), for the TRITC channel the laser was set at 0.9 (HV = 40).

For imaging of transiently produced ACTB mRNA and stress granules in fixed samples, the pinhole size was 63.9 μm, the power of the 488 nm laser line was 1.5 (80 V gain), the power of the 561 nm laser line was 2.0 (110 V gain) and the power of the 636 nm laser line was 6.0 (110 V gain). For imaging of endogenous ACTB mRNA, the pinhole was 58.7 μm, the laser power of the 488 nm laser line was 0.4 (40 V gain) and the laser power of the 638 nm laser line was 12 (120 V gain). The signal from the 638 nm laser line was integrated 16 times.

For live imaging of HeLa cells, the pinhole was 51.1 μm, the laser power for the 561 nm laser line was 2-5 with a gain of 50-80 V (adjusted for optimal contrast to account for differences in bead loading efficiency) and the laser power for the 488 nm laser line was 0.5 (50 V gain).For live imaging of GFP-SMN with or without co-transfection of a plasmid to produce A_T_-U1 RNA, the pinhole was 51.1 μm, the laser power for the 561 nm laser line was 2 (50 V gain) and the laser power for the 488 nm laser was 0.5. The gain was adjusted for optimal signal without over-exposure of puncta in the green channel (30 V gain for the single GFP-SMN cells shown in Supplementary Fig. 32 and 50 V gain for the double transfected cell shown in Fig. 5c). For imaging of endogenous SMN and DDX20 simultaneous with U1 snRNA (either endogenous U1 snRNA or transfected to produce A_T_-U1 snRNA) in fixed HeLa cells, the pinhole was 49.8 μm, the laser power for the 405 nm laser line was 2.0 (90 V gain), the laser power for the 561 nm laser line was 0.3 for SMN imaging and 1.0 for DDX20 imaging (40 V gain in all cases) and the laser power for the 638 nm laser line was 10.0 (120 V gain). For imaging of transiently co-transfected GFP-SMN and A_T_-tagged U1 in fixed HeLa cells, the pinhole was 49.8 μm, the laser power for the 488 nm laser line was 0.3 (25 V gain) and the laser power for the 638 nm laser line was 10.0 (120 V gain). For imaging of coilin colocalization in fixed HeLa cells, the pinhole was 0.8 AU, the laser power for the 405 nm laser line was 1.0 (100 V gain), the laser power for the 488 nm laser line was 2.0 (40 V gain) and the laser power for the 638 nm laser line was 4.0 (110 V gain).

For live imaging of Broccoli-tagged 5*S* and U6 RNA, published protocols were used^26^.Briefly, plasmids pAVU6+27-F30-2xdBroccoli and pAV5S-F30-2xdBroccoli^26^ were transfected in HEK293T cells and split into imaging dishes 48 h post transfection. 24 h later, DFHBI-1T was added at a final concentration of 40 μM and cells were imaged under widefield illumination conditions as recommended^26^ using a 60x oil objective and 250 ms exposure. No ND filters were used.

### Immunofluorescence

Cells were fixed in 4% paraformaldehyde (EM grade, Electron Microscopy Sciences) for 10 min and slides were rinse three times in PBS and permeabilized for 10 min in PBS / 0.2% Triton X-100 at room temperature, followed by three rinses in PBS. After blocking for 30 min at room temperature in PBS / 5% FBS, slides were incubated with the primary antibody against DDX20 or SMN (both 1:200 dilution) in PBS / 5% FBS at 4°C overnight. The DDX20 antibody was a mouse monoclonal antibody (sc-57007, Santa Cruz Biotechnology) and the SMN antibody was a rabbit polyclonal antibody (sc-15320, Santa Cruz Biotechnology). After three rinses in PBS, slides were incubated with the secondary antibody (1:1,000 dilution in both cases) and Hoechst nuclear dye (1:10,000 dilution) for 90 min at room temperature. The secondary antibody for DDX20 was a goat anti-mouse Alexa 594 antibody (Invitrogen) and the secondary antibody for SMN was a donkey anti-rabbit Alexa 568 antibody (Invitrogen). Slides were rinsed three times in PBS and once in water. If no FISH was performed subsequently, slides were mounted.

### Fluorescence in situ hybridization (FISH)

When no IF was performed prior to FISH, cells were fixed in 4% paraformaldehyde and permeabilized in 0.2% Triton X-100 as described above. When FISH was performed after IF, no additional permeabilization step was performed. Cells were dehydrated sequentially in 70%, 95% and 100% ethanol for 5 minutes each. After a two minute drying step, cells were rehydrated in 2x SSC (1x SSC is 150 mM NaCl, 15 mM sodium citrate dihydrate, pH 7.0) / 50% formamide (molecular biology grade) for 5 min. Slides were pre-hydrated in pre-hybridization solution for 30 min at 37°C. The pre-hybridization solution contained 50% formamide, 2x SSC,0.5 mg/mL UltraPure Salmon sperm DNA (ThermoFisher Scientific), 1 mg/mL UltraPure BSA (ThermoFisher Scientific), 0.13 mg/mL *E. coli* tRNA (Sigma Aldrich), 1 mM Vanadyl ribonucleoside complexes solution (Signal Aldrich) and 100 mg/mL dextran sulfate in ultrapure water. After pre-hybridization, samples were hybridized with the probe in pre-hybridization solution (see Supplementary Table 10 for properties) overnight at 37°C. On the following day, samples were washed twice in 2x SSC / 50% formamide for 30 minutes each. Slides were rinsed in PBS once and sealed.

### Northern blotting

The production and processing of aptamer fusion variants was assessed in 293T cells. Prior to seeding, dishes were coated with 10 μg/mL Poly-L-Lysine for 1 h and rinsed. For each aptamer variant, ~2×10^6^ cells were seeded in a 10 cm dish. On the following day, 15 μg plasmid DNA was transfected per 10 cm dish using the *Trans*IT (Mirus) chemical transfection system according to the manufacturer’s recommendations. Cells were harvested 48 h after transfection and cell pellets were frozen at -80°C.

The total RNA was extracted using the RNeasy (Qiagen) kit according to the manufacturer’s recommendation. Briefly, cell pellets were thawed on ice, resuspended in buffer RLT and lysed by passing six times through a syringe needle. The lysate was supplemented with 70% ethanol and applied to an RNeasy spin column. After on-column DNase treatment and washing steps, RNA was eluted in water. The final RNA concentration was typically 500-1000 ng/μL. The RNA was stored at -80°C for no longer than one week before proceeding.

All solutions for gel electrophoresis and Northern blotting were made in diethylpyrocarbonate (DEPC)-treated and autoclaved water to ensure removal of RNase. For each blot, RNA samples were normalized for total RNA amount (10 μg total RNA per lane).Samples with the desired amount of total RNA were tried in a SpeedVac and the RNA was brought up in 15 μL of RNA sample buffer (50% v/v formamide, 6.3% v/v formaldehyde, 0.2M MOPS, 50 mM sodium acetate pH 5.2, 10 mM EDTA pH 8.0) plus 2 μL of RNA loading buffer (50% glycerol, 1 mM EDTA pH 8.0, 0.4% Bromophenol blue, 1 mg/mL ethidium bromide). The samples were heated at 65°C for 5 min and then loaded on a 1% agarose/formaldehyde gel (50 mM sodium acetate pH 5.2, 10 mM EDTA pH 8.0, 1% w/v agarose, 6.3% formaldehyde). 10 μL of a Low Range ssRNA Ladder (New England Biolabs) was loaded as well. The gel was run in running buffer (50 mM sodium acetate pH 5.2, 10 mM EDTA pH 8.0, 6.3% formaldehyde) at 60 V for 2.5 h. To assess the quality of the isolated RNA, the gel was then stained with ethidium bromide (10 μL of 10 mg/mL ethidium bromide in 400 mL running buffer).

After destaining in water for 5 min, the RNA was transferred to a nylon membrane as follows. Immersed in 10X SSC, two pieces of Whatman paper were placed on a stack of blotting paper, followed by the agarose gel and the pre-wet membrane (Amersham Hybond-N^+^, GE Healthcare). The membrane was covered with two pieces of pre-wet Whatman paper and the RNA was allowed to transfer to the membrane by capillary transfer overnight. The RNA was then crosslinked to the membrane by exposure to UV light, followed by a washing step in 0.1X SSC and 0.1% sodium dodecyl sulfate (SDS) at 65°C for 1 h. The membrane was prehybridized for 1 h in hybridization solution (6X SSC, 10X Denhardts solution (Life Technologies), 0.1% SDS) at 42°C. The membrane was then incubated with hybridization solution supplemented with the radioactive, labeled probe overnight at 42°C. After 3 wash steps for 5 min at room temperature in washing solution (6X SSC, 0.1% SDS) and an additional wash step at 48°C for 20 min, the membrane was tried and exposed overnight. Exposure and visualization of the ^32^P signal was done using the Phosphor Screen and Cassette System from GE Healthcare. The membrane was stripped by repeated microwaving in 0.1X SSC, 0.1% SDS to subsequently hybridize with another probe.

Radioactive probes were produced as follows. The sequence of each probe is indicated in Table S5 and probes were purchased as DNA primers from IDT. From a 100 μM stock of this DNA primer, 2 μL were mixed with 2 μL 32P-ATP, 2 μL T4 Polynucleotide Kinase (PNK) (NEB), 2 μL of 10X T4 PNK buffer (NEB) in water and incubated at 37°C for at least 1 h. The DNA was then passed over a G-25 column (GE Healthcare) according to the manufacturer’s recommendations to remove unincorporated ATP.

### Data availability statement

The datasets generated during and/or analyzed during the current study are available from the corresponding author on reasonable request.

